# Monitoring fish spawning sites in freshwater ecosystems using low-cost UAV data: A case study of salmonids in lakes in Iceland

**DOI:** 10.1101/2021.06.12.448199

**Authors:** Lieke Ponsioen, Kalina H. Kapralova, Fredrik Holm, Silvia García Martínez, Benjamin D. Hennig

**Affiliations:** Institute of Life and Environmental Sciences, University of Iceland, Askja – Sturlugötu 7, 102 Reykjavík, Iceland

## Abstract

Low-cost unmanned aerial vehicles (UAV), widely known as drones, have become ubiquitous and improved considerably in their technical capabilities and data quality. This opens new opportunities for their utilisation in scientific research that can help to reduce equipment and data collection costs. Remote sensing methods in ecological fieldwork can be a suitable approach to complementing, augmenting or even replacing certain aspects of fieldwork. In this study we tested the suitability of UAV for the detection of salmonid spawning grounds in two lakes in Iceland, Thingvallavatn and Ellidavatn. Salmonids are very susceptible to environmental changes, especially during embryonic development when highly oxygenated water flow and low temperatures are required. Monitoring the changes of the redd density over time will help understand the population dynamics of salmonid species and create strategies for species conservation. As part of this pilot study, we conducted aerial surveys during the spawning seasons in both locations in 2018 recording standard photographs in the visible spectrum (red, green, blue) to fully cover the respective areas of interest. Different flight altitudes were recorded to test the effects of image resolution on the final analyses. The images were then processed by applying standard remote sensing analyses in the software ENVI. The maximum likelihood classification combined with post-classification improvement methods resulted in satisfactory accuracies that are valuable for further monitoring efforts. From these findings we established a workflow that allows the implementation of UAV in ecological fieldwork for regular long-term observations. We discuss our experiences with regards to their utility, their limitations and identify future directions of research for implementing the potential that low-cost approaches have in supporting ecological studies of freshwater ecosystems.

## Introduction

Salmonids are cold temperate species that are expected to be negatively impacted by the warming of aquatic habitats and the correlated oxygen depletion due to climate change (Jankowski et al., 2006; Jonsson & Jonsson, 2009). Because of their inability to shift their geographical range to better environmental conditions, landlocked populations and populations at the southern end of their distribution range are especially vulnerable. Rising temperatures often lead to increased physiological stress and depletion of salmonids’ energy reserves (Jonsson & Jonsson, 2009), increased susceptibility and exposure to diseases (Karvonen et al., 2010) and disruptions to their breeding efforts (Pankhurst & King, 2010). Highly oxygenated water flow and low temperatures are required for salmonids’ successful embryonic development, making them especially vulnerable during this time (Moyle, 2002). However, the majority of research and monitoring efforts are usually focussed on later life stages (Binder et al., 2018; Hall & Werner, 1977; Hankin & Reeves, 1988; Keenleyside, 1962; Leander et al., 2019; Riley et al., 1992; Satterthwaite et al., 2012; Schill & Griffith, 1984; Vélez-Espino et al., 2016; Watson et al., 2019; Wilson et al., 2004). Evidence is showing that our current climate is changing more rapidly than before (Ficke et al., 2007). Due to their long generation time (Willson, 1997), the monitoring of salmonid populations during their spawning and subsequent embryonic development of their offspring would provide information on the effects of climate change much faster than the data that would be otherwise gathered on later life stages.

Most species belonging to the *Salmonidae* family are characterised as gravel nests spawners (Nika et al., 2011). Due to their appearance as irregular or regular shapes that contrast with the undisturbed area surrounding them (Bjornn & Reiser, 1991), the spawning redds (*i*.*e*. nests) are visible from the air. This makes monitoring of the spawning redds from the shore and directly above the water possible. Although redd count data are an important source of information for management purposes, including monitoring population size and estimating carrying capacity of spawning habitats (Rieman & Mcintyre, 1996), redd estimates are predominantly done by manual counting from the shoreline. This method is often criticised for the common occurrence of sampling errors resulting in an underestimation of redd numbers (Dauphin et al., 2010; Murdoch et al., 2018). Sampling error can be assigned to a wide variety of factors, for example low visibility due to physical characteristics of the spawning redd location (*e*.*g*. water depth), incomplete sampling of spawning areas, and inexperienced observers, among others (Dauphin et al., 2010; Dunham et al., 2001). Apart from sampling errors, manual redd counting is a time-consuming effort, it does not allow for precise quantification of redd changes over time, and it is usually performed by different observers that use dissimilar methodologies on different temporal and/or spatial scales. Due to the inherent high variability of the method and the inability to compare results, the shortcomings of the manual redd count make monitoring using this method rather difficult (Schuett-Hames et al., 1996).

The use of unmanned aerial vehicles (UAV), also known as drones, for counting salmonid redds is being explored as an answer to these shortcomings (Groves et al., 2016). Groves et al. (2016) used an UAV to obtain video footage as an alternative to manned helicopter flights to visually identify Chinook salmon (*Oncorhynchus tshawytscha*) redds in the Lower Snake River, USA. They found UAV-based counting to be more accurate than redd counts done from a helicopter flight. While this method only partially replaced manual redd count by remote sensing, other methods for identifying salmonid redds have explored the feasibility of completely replacing manual count with aerial surveys in combination with semi-automated remote sensing approaches. For example, Roncoroni & Lane (2019) used UAV in combination with Structure from Motion (SfM) photogrammetry and digital elevation models to detect brown trout (*Salmo trutta*) redds and reported that the use of this approach helped identify more redds than using visual detection. Another study by Harrison et al. (2020) applied machine learning and object-based classification techniques to UAV-based imagery to assess the use of both RGB and hyperspectral imagery. They found that both sources of imagery could be used to identify redds, but with varying degrees of accuracy. Despite the current development of techniques that use remote sensing to detect redds, the application of such techniques is not yet an integral part of large-scale monitoring programmes.

The implementation of UAV in monitoring programmes requires the establishment of routines and procedures that can easily be applied and adapted to specific project needs. The cost of such efforts must also be taken into account in order to keep monitoring efforts manageable and effective. This gives low-cost UAV an advantage, and provides further benefits due to their quick and relatively easy data collection (Calvario et al., 2017). The goal of this study was to develop an easy, accessible and low-cost method to map salmonid spawning redds. We created a pipeline for detecting salmonid redds using a semi-automated approach by applying a pixel-based classification technique to UAV derived imagery. In this study, we present the pipeline developed to detect salmonid spawning grounds which was tested at two different Arctic charr (*Salvelinus alpinus*) spawning grounds in Iceland.

## Material and methods

### Study areas

Two study areas (*i*.*e*. Thingvallavatn, Ellidavatn) with contrasting environmental characteristics (Table 1) were selected among Icelandic freshwater ecosystems to establish and validate the method presented here.

**Table 1.** Summary of environmental conditions in Thingvallavatn and Ellidavatn. From left to right: the spawning site, the date and time of the aerial survey, the contrast between spawning redds and other environmental features, the topography of the spawning area, reflectance similarities of the spawning redds and other topographical features, and sunlight reflection.

| Site | Date and time | Contrast | Topography | Reflectance similarities | Light |
| --- | --- | --- | --- | --- | --- |
| Thingvallavatn | July, 19:30 | High | Heterogeneous | Low | High |
| Ellidavatn | October, 12:00 | Low | Homogeneous | High | Low |

The method was first established in lake Thingvallavatn, located in southwestern Iceland. The lake has a surface area of 83 km^2^ and a maximum depth of 114 m, making it the largest and deepest natural lake of Iceland (Adalsteinsson et al., 1992). Two salmonid species reside in the lake, Arctic charr (*Salvelinus alpinus*) and brown trout. Four morphs of Arctic charr are found in the lake which differ in life history characteristics, morphology, genetics, and other characteristics. The morphs can be classified into two morphotypes depending on which habitat they occupy: benthic (*i*.*e*. small benthic and large benthic), and pelagic (*i*.*e*. planktivorous and piscivorous) (Snorrason et al., 1989). In this study we focussed on the well-studied spawning grounds of Ólafsdráttur, an area of the lake which is known to host the spawning of the large benthic Arctic charr in July and August each year. Ólafsdráttur is located on the northeastern shore of the lake within the protected Thingvellir national park and is characterised by the influx of cold water from springs, which makes for a stable, cold temperature (3.2-3.8 °C) throughout the year (Adalsteinsson et al., 1992; Ólafsson, 1992). The spawning redds in this area are located in both very shallow (*i*.*e*., 0.5-1.5 m) and deeper water (*i*.*e*., 1.5-5 m).

The second study area, lake Helluvatn connected to lake Ellidavatn, is situated in the vicinity of the urban area of the capital city of Iceland, Reykjavík. Ellidavatn is a small and shallow lake with a surface area of 2 km^2^ and 1 to 3 m depth (Björnsson, 2001). The two salmonid species, Arctic charr and brown trout, found in Thingvallavatn also inhabit Ellidavatn and Helluvatn (Malmquist et al., 2009). Arctic charr primarily spawn in the lake from September until November (Björnsson, 2001; Malmquist et al., 2009). In the last years, an expansion of the city led to new residential areas being built closer to the lake’s nature reserve. This has led to increased human impact due to street and drain water from the residential area flowing directly into the rivers surrounding the lake (Guðmundsdóttir, 2012).

This study is based on the assumption that salmonid redd structures in shallow water areas can be distinguished by their visible reflection which makes the use of standard UAV a viable option. Due to spawning Arctic charr females cleaning the rocks from debris, silt, and algae to prepare the spots for spawning, redds can be identified from the air as dark areas of gravel and rocks (Figure 2D). The distinction of spawning redds and other geographical features depends on the surrounding environment. The two study areas chosen in this study represent contrasting environments (Table 1). Thingvallavatn is characterised by variance in deep water level, while in Ellidavatn the deep-water level presents a more homogeneous ground level. Furthermore, the occurrence of water vegetation differs between areas with vegetation emerging at similar ground level as the spawning redds in Ellidavatn making their distinction more challenging.

The study was realised in four main stages which are described below: The initial data acquisition at the first case study area was followed by the main data processing steps which were then concluded with an accuracy assessment to evaluate the quality of the results (Figure 1). Once a successful workflow had been successfully established, the procedure was transferred to our second case study area for validating the method.

**Figure 1.**
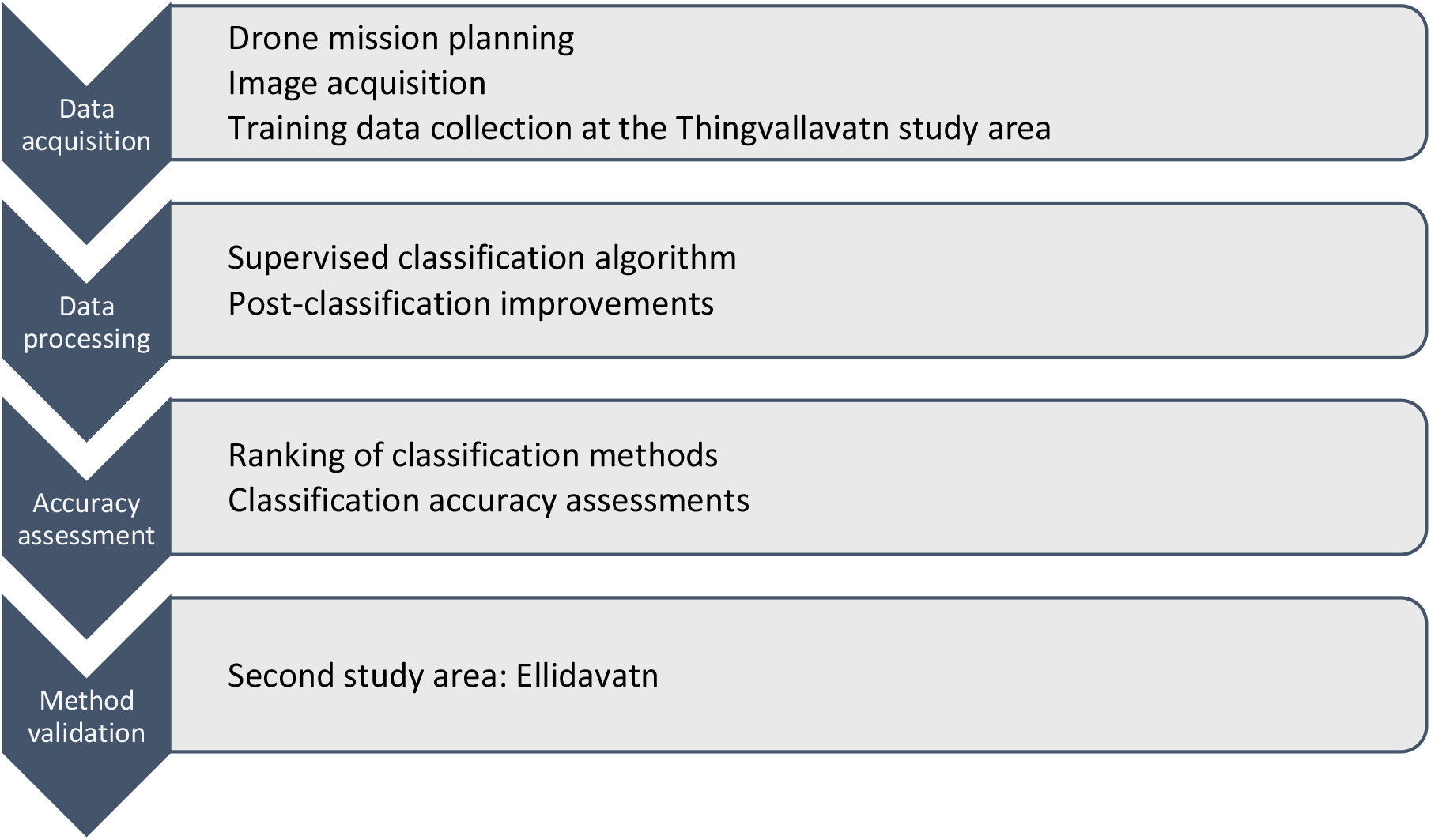
Main steps of establishing the method to detect spawning grounds.

### Data acquisition

A certificate of authorisation was obtained from National Park Thingvellir for the UAV survey in Thingvallavatn. Data collection was undertaken using a DJI Mavic Pro drone (Table S1) equipped with a true colour camera and a remote controller to command the drone. The remote controller was operated on an Android smartphone with the DJI GO 4 application developed by the UAV manufacturer. To avoid the sun’s reflection on the water surface, the aerial surveys were taken before sunset, and minimal wind was necessary to avoid ripples in the water surface. The aerial surveys were completed on July 27th 2018, between 17:00h and 19:00h in Thingvallavatn, and on October 31st 2018, between 12:00h and 12:30h in Ellidavatn. The timing of the survey was chosen in consideration with the spawning season of the Arctic charr and optimal weather conditions. Recorded weather conditions in the Thingvallavatn survey area were fair weather with few clouds, a total of 18.26 hours of sunlight, and 2 m/s wind SSE direction. In Ellidavatn wind speed was at 3 m/s from E direction during the survey. The total duration of the Thingvallavatn survey was one full hour (three full batteries), narrowing the aerial time but obtaining sufficient coverage of the study area. For Ellidavatn half the time was needed due to a smaller area covered by the sampling site. The flight direction of the surveys was controlled manually by a professional drone pilot. It followed the shallow shoreline (0-10 m) where the spawning redds were more concentrated. The flight path at Thingvallavatn was taken over the same areal extent at 10 m, 50 m, and 100 m with the intention of capturing different resolution images for comparison.

The challenges of this methodology lie in the heterogeneous structure of the spawning redds and the similar reflectance with water vegetation and deep water that can interfere in the classification of the features. To establish reliable ground truth information for each endmember, specialists in Arctic charr and field observations from the study area helped identifying the spawning grounds on the aerial images. Fifteen training samples of each spectral class were selected to allow for reasonable estimates to determine the mean vector and the covariance matrix for the maximum likelihood classification (Richards & Jia, 1999).

### Data processing

The software ENVI version 5.1 (Exelis Visual Information Solutions, Boulder, Colorado) was used for processing and classification analysis. For data processing, only images with the least sunlight reflection and wind ripples were selected. Eight supervised image classification methods were tested for their accuracy at classifying spawning redds (García Martínez, 2019). Considering that the maximum likelihood classification had the highest overall accuracy when classifying spawning redds, it was decided that this approach would be taken forward for further analyses in this study to improve the results specifically for the redds feature class. The maximum likelihood classification method assigns individual pixels to the class with the highest *a posteriori* probability under the assumption of a normal distribution (Richards & Jia, 1999). When using this method in ENVI, no probability threshold was set to classify all pixels in the image.

Three post-classification methods were applied in order to improve the accuracy by correcting isolated or misclassified pixels. First, the classified output was filtered by removing spurious pixels with a majority-minority analysis (Gurney & Townshend, 1983). This analysis changes spurious or “false” pixels to the class value that the majority of the pixels in the manually indicated kernel belong to. The following parameters were selected: majority for the analysis method, a kernel size of 3, and a centre pixel weight of 1. In addition, isolated pixels were corrected using the sieve classes method (Exelis Visual Information Solutions, 2009; Su, 2016). This method looks at neighbouring pixels to determine if a pixel is grouped with the same endmember classes surrounding the pixel. For this study pixel connectivity was set to four and the minimum size to two. Lastly, the clump classes method was applied to clump similarly classified areas together adjacent from each other (Su, 2016). Following a visual examination of the initial classification results, the size parameter for this method was set to three.

Overall accuracy of the applied supervised algorithm is expressed as the percentage of correctly classified pixels of all endmember classes. Producer’s accuracy (PA) is defined as the probability that each endmember class is classified correctly. User’s accuracy (UA) is defined as the probability that the classification map represents the ground truth data. Furthermore, the kappa coefficient was used to evaluate the classification accuracy and can be interpreted as a value ranging from 0 to 1 that explains the difference between the observed classification of the endmember classes and the reference data (Campbell, 2002).

## Results

The method to classify spawning redds from UAV imagery was established using the well-studied Ólafsdráttur spawning grounds in Thingvallavatn where the contrast of the spawning redds was high and reflectance similarities were low. Aerial images were taken at altitudes of 10 m, 50 m, and 100 m above water surface level (Figure 2A-C). After these three different altitudes were tested for endmember collection and utilising the algorithms, 10 m and 100 m imagery were discarded. The 10 m imagery was discarded due to an insufficient number of classes and the 100 m imagery due to reflectance and low accuracy results. The 50 m imagery was deemed suitable for the classification methods after a visual examination, as these images provided the optimal contrast and lowest environmental reflectance in relation to the spawning grounds as the main features of interest in this analysis.

**Figure 2.**
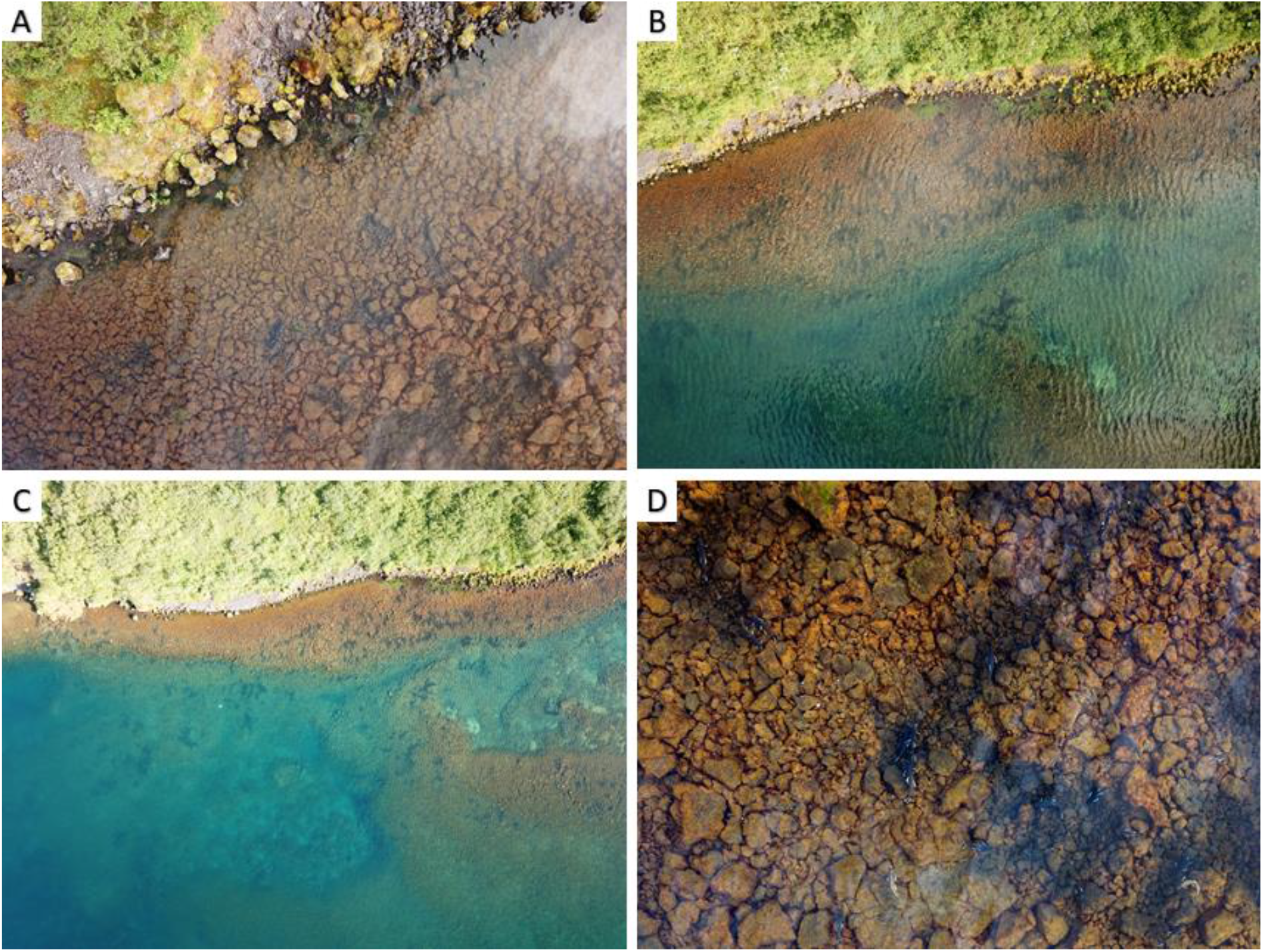
A-C: Aerial images of Ólafsdráttur spawning grounds at different altitudes: 10m (A), 50m (B), and 100m (C). D: close-up view of the dark spawning redds that are made by salmonids.

Using the maximum likelihood classification method with the 50 m image of the spawning grounds, fifteen training areas were selected for each spectral class with an average of 120 pixels each, to not over- or undertrain the supervised classification method. The accuracy assessment reported a producer’s accuracy of 88.62% and a user’s accuracy of 94.86% for classifying spawning redds before any post-classification improvements were applied. Six endmember classes were obtained after applying the maximum likelihood classification algorithm and post-classification improvements (*i*.*e*. majority/minority analysis, sieve classes, clump classes) to remove any false or spurious pixels (Figure 3).

**Figure 3.**
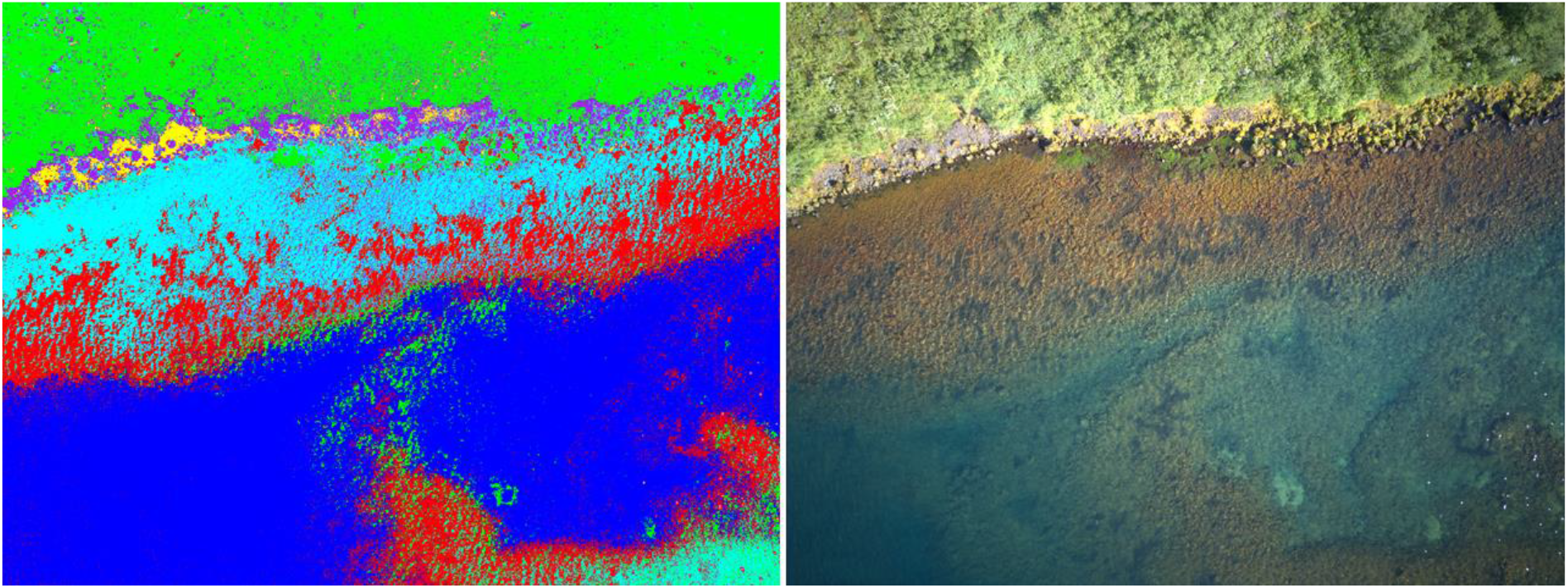
Overview of the endmember classes obtained with the maximum likelihood classification in Ólafsdráttur. Classified pixels as spawning redds are in red, underwater rocks in cyan, vegetation in green, deep water in blue, shoreline in yellow, and surface rocks in purple.

Class 1 (spawning redds; in red) is a dominant class covering parts of the shallow area, close to the shore, and in the centre of the image in the deeper waters. This class can be described as a number of small, irregular groups of pixels connected by few pixels surrounding the groups. The UAV image shows low reflectance by a dark colour at places where this class is dominating. Class 2 (underwater rocks; in cyan) is distributed covering the pixels adjacent to class 1 and completes the shallow area. The major difference between these two classes is the lack of irregular groups of pixels in class 2. The UAV image shows a lighter colour than class 1. Class 3 (vegetation; in green) can be recognised as a green coloured area located on land with few pixels close to the shoreline in the UAV image. This class covers the entire eastern part of the image in one group of pixels. Class 4 (deep water; in blue) can be described as a very defined and smooth polygon class, located in deeper water. The UAV image shows a deeper, darker area with few ripples caused by wind on the water surface. Class 5 (shoreline; in yellow) is defined by little groups of pixels adjacent to class 3, located over the shoreline. This class is very heterogeneous and has few pixels per group that do not cover a large area. The UAV image shows very light areas where this class covers. Lastly, class 6 (surface rocks; in purple) occurs over the shoreline and in some parts in shallow water. The UAV image shows high reflectance from sunlight.

An accuracy assessment was performed using the selected training data and test data to estimate the quality of the classification results using the maximum likelihood classification method after post-classification improvements were applied (Table 2). The classification and the ground truth data were combined in an error matrix to estimate the number of correctly classified pixels. Most importantly, the producer’s accuracy for classifying spawning redds was 90.68% and 97.07% for the user’s accuracy. While the spawning redds were mostly correctly classified, 26 pixels were classified as underwater rocks, 8 pixels as shoreline, 5 pixels as deep water, and 1 pixel as surface rocks.

**Table 2.** Error matrix of the maximum likelihood classification in pixels for Thingvallavatn. PA and UA are reported after applying post-classification methods.

| Class | Vegetation | Shoreline | Surface rocks | Deep water | Underwater rocks | Spawning redds | Total | User's accuracy (%) |
| --- | --- | --- | --- | --- | --- | --- | --- | --- |
| Unclassified | 0 | 0 | 0 | 0 | 0 | 0 | 0 |  |
| Vegetation | **2485** | 0 | 8 | 0 | 0 | 0 | 2493 | 99.68 |
| Shoreline | 0 | **1312** | 25 | 0 | 0 | 0 | 1337 | 98.13 |
| Surface rocks | 12 | 216 | **1506** | 0 | 51 | 0 | 1785 | 84.37 |
| Deep water | 0 | 0 | 0 | **2396** | 0 | 111 | 2507 | 95.57 |
| Underwater rocks | 0 | 1 | 214 | 0 | **921** | 25 | 1161 | 79.33 |
| Spawning redds | 0 | 8 | 1 | 5 | 26 | **1323** | 1363 | 97.07 |
| Total | 2497 | 1537 | 1754 | 2401 | 998 | 1459 | 10646 |  |
| Producer's accuracy (%) | 99.52 | 85.36 | 85.86 | 99.79 | 92.28 | 90.68 |  |  |

The above results are all reported after applying post-classification methods. The addition of these classification improvements raised the overall accuracy of the maximum likelihood classification from 91.27% to 93.40% (Table S2).

The Ólafsdráttur spawning grounds had high contrast of spawning redds and low reflectance similarities of geographical features, so this method was applied in an environment with very different conditions, such as the Ellidavatn spawning grounds with low contrast and high reflectance similarities. Six endmember classes were obtained after applying the maximum likelihood classification algorithms (Figure 4). Class 1 (spawning redds; in red) appears in the shallow water and occurs as irregular groups of pixels with connected by few pixels surrounding the groups. Additionally, few irregular groups are located on land. The UAV image shows a dark colour where the pixels are located. Class 2 (underwater rocks; in cyan) is a dominant class covering the shallow area. This class is adjacent to class 1 and 4, and is characterised with a lighter colour in the UAV image. Class 3 (vegetation; in green) is a dominant cover class in the land area and has a high density of groups of pixels connected to each other. In the UAV image this group can be recognised due to its light colouring. Class 4 (aquatic vegetation; in yellow) covers areas close to the shore and further away towards deeper water. The groups of pixels are of irregular shape and are found in high-density. The UAV image shows a similar colour reflection to class 1, but differences are found in the density and location of the class. Furthermore, some pixels are located in a small shaded part of the land area. Class 5 (ice; in blue) consists out of a high-density main group of pixels located in the North-western edge of the image with an irregular shape and furthermore, few pixels distributed not larger than a few pixels together. The UAV image shows that this class is fully distributed on land. Lastly, class 6 (human structure; in white) is very heterogeneous and does not cover a large area. The class has a main group of pixels in a square shape and some more pixels spread out in small clusters consisting of only a few pixels. The UAV image shows this class located on land recognised by a grey colour.

**Figure 4.**
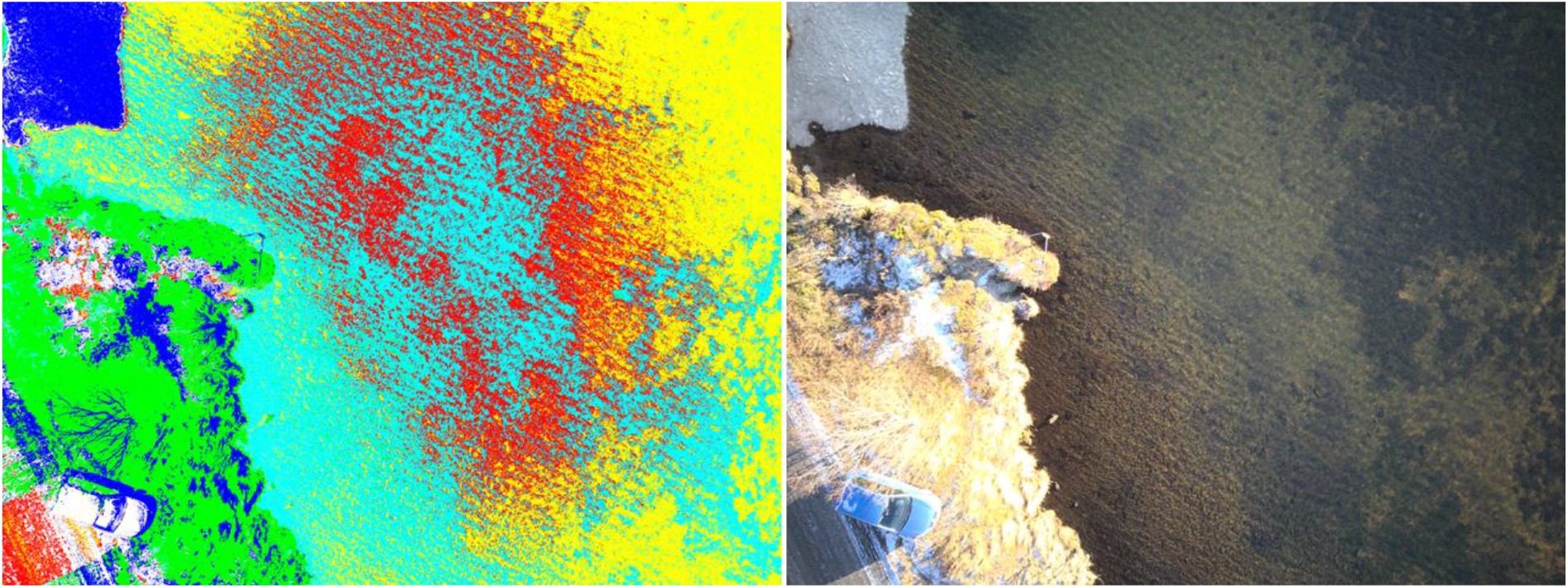
Overview of the endmember classes obtained with the maximum likelihood classification in Ellidavatn. Classified pixels as spawning redds are in red, underwater rocks in cyan, vegetation in green, aquatic vegetation in yellow, ice in blue, and human structures in white.

An accuracy assessment was performed to estimate the quality of the classification results using the maximum likelihood classification method (Table 3). The classification and the ground truth data were combined in an error matrix to estimate the amount of correctly classified pixels. The producer’s accuracy for classifying spawning redds reported 83.41% and 58.83% for the user’s accuracy. Misclassified pixels were classified as either human structures, underwater rocks, or aquatic vegetation.

**Table 3.** Error matrix of the maximum likelihood classification in pixels for Ellidavatn. PA and UA are reported after applying post-classification methods.

| Class | Ice | Vegetation | Aquatic vegetation | Underwater rocks | Human structure | Spawning redds | Total | User's accuracy (%) |
| --- | --- | --- | --- | --- | --- | --- | --- | --- |
| Unclassified | 0 | 0 | 0 | 0 | 0 | 0 | 0 |  |
| Ice | 634 | 0 | 0 | 0 | 0 | 0 | 634 | 100.00 |
| Vegetation | 10 | 865 | 0 | 0 | 48 | 0 | 923 | 93.72 |
| Aquatic vegetation | 0 | 0 | 495 | 22 | 0 | 11 | 528 | 93.75 |
| Underwater rocks | 0 | 2 | 3 | 613 | 1 | 87 | 706 | 86.83 |
| Human structure | 0 | 0 | 0 | 3 | 415 | 6 | 424 | 97.88 |
| Spawning redds | 0 | 0 | 48 | 136 | 182 | 523 | 889 | 58.83 |
| Total | 644 | 867 | 546 | 774 | 646 | 627 | 4104 |  |
| Producer's accuracy (%) | 98.45 | 99.77 | 90.66 | 79.20 | 64.24 | 83.41 |  |  |

To obtain the best results using the maximum likelihood classification method the post-classification methods, majority-minority analysis, sieve classes method and clump classes, were applied after classifying the pixels. The addition of these classification improvements raised the overall accuracy of the maximum likelihood classification from 82.48% to 86.40% (Table S2).

## Discussion

This study aimed at developing an easy, accessible and low-cost method for mapping salmonid spawning grounds in shallow freshwater areas. The analysis of UAV-derived imagery in the contrasting case study areas situated in the subarctic region showed a successful application of a pixel-based classification method that was able to identify spawning redds from RGB imagery with high accuracy. Imagery taken from a height of 50 m was deemed suitable for classifying spawning redds after which fifteen training areas were selected for each spectral class with an average of 120 pixels each. The maximum likelihood classification method was run with resulting high accuracies for producer’s and user’s accuracy. By correcting isolated or misclassified pixels three post-classification improvement methods were applied to improve accuracy: majority-minority analysis, sieve classes method and clump classes.

The use of remote sensing for mapping and monitoring spawning grounds provides improvement over traditional means of monitoring due to the method being safer (Groves et al., 2016), more accurate, and more time and cost efficient (Harris et al., 2019; Harrison et al., 2020). However, determining a suitable UAV and analysing platforms can be difficult due to the many options that are available with varying levels in cost and difficulty (Harris et al., 2019). Our approach makes use of an off-the-shelf UAV equipped with a standard RGB camera which significantly reduces the cost of the UAV since no specialised equipment is necessary (Harris et al., 2019). Hyperspectral cameras have also shown to successfully detect spawning redds (Harrison et al., 2020) but they come with considerably higher costs and often lower spatial resolution than RGB (Hardin et al., 2019). Our approach has shown the effectiveness of using RGB imagery to map spawning redds using a semi-automated approach in two contrasting environments. Analysing UAV derived imagery using semi-automated approaches would provide an advantage over manual counting (Groves et al., 2016) when large quantities of images need to be analysed in case of frequent monitoring. Semi-automated approaches reduce analysing time and improve accuracy in case of observer bias which is often problematic when manual counts are concerned (Dunham et al., 2001). Furthermore, the combination of a single RGB image and the maximum likelihood classification method makes our method accessible to a wide range of users that intent to map and monitor features such as nesting sites in the shallower parts of lakes. Needing only a single image for a successful analysis reduces the time and effort during data collection, as opposed to the approach of Roncoroni and Lane (2019) where multiple images are required to extract elevation data. Harrison et al. (2020) noted that RGB is more easily obtainable than hyperspectral imagery and would make monitoring on frequent basis more accessible. Furthermore, processing of this type of data is relatively easy and quick to achieve once a workflow has been established. Moreover, our method has the potential to be used on composite images that encompass entire spawning grounds without expecting considerable setbacks.

Large-scale monitoring programs are needed to assess ecosystem changes caused by climate change and other anthropogenic stressors. Our proposed method allows for regular observation of changes in spawning sites with minimal effort. Climate change has been warming freshwater ecosystems at higher latitudes and altitudes disproportionally (Hassan et al., 2005). Furthermore, other anthropogenic stressors have negatively affected spawning grounds in freshwater ecosystems (Dudgeon et al., 2006). Excessive accumulation of fine sediments in spawning gravels that originates from land uses that increase sediment yields, such as logging and road construction (Everest et al., 1987) as well as rising temperatures have negative impacts on salmonid embryo development and fry emergence (Heywood & Walling, 2007; Jeuthe et al., 2016; Ojanguren et al., 1999; Suttle et al., 2004). Other anthropogenic stressors lead to the destruction of spawning habitats, for example through dredging lake bottoms or exposing them to intolerable fluctuations of the water level (Maitland, 1995), and the construction of dams that make spawning grounds inaccessible for anadromous salmonids (Quiñones et al., 2015). In the case of our study system Thingvallavatn, a small dam was built in 1959 on the southern outlet of the lake (Sturlaugsson & Malmquist, 2011). The outflow of the lake had to be directed through a short tunnel leaving the former outlet mostly dry. The installation of the dam and the tunnel had a significant impact on the brown trout population of the lake. It is believed that a major spawning site at the outlet was spoiled and the drying-up of the former outlet ruined an extremely productive community of blackfly larvae (*Simulium vittatum*) that formed optimal feeding grounds for juvenile trout (Malmquist, 2011). These however are only speculations since the massive decline of the brown trout population came to light more than 20 years later when in the 1980s during a lake-wide survey only a handful of trout specimen were caught (Sandlund et al., 1987). These are just a few examples on how human impact has negatively influenced spawning habitats and while in some places action has been undertaken to, for example, remove dams to mitigate these effects, the removal is unlikely to restore these ecosystems the way they were previously since the negative effects of the dam have been amplified through time (Quiñones et al., 2015). Often these negative impacts are only observed years later as in the case of the decline of brown trout in Thingvallavatn. Systematic monitoring of the shorelines of the lake through a standardised monitoring system could have shown the effects of stressors much sooner.

Our approach to detect spawning grounds has the potential for future use in complementing fieldwork that aims at monitoring changes in salmonid spawning grounds in different environmental conditions. Scaling this method to other freshwater ecosystems would allow to assess the full potential of this method in detecting spawning redds from a range of different species. We show the successful application of this method in two very contrasting lake environments which indicates the promising potential of this low-cost method.

Remote sensing is a helpful tool for monitoring from small to large spatial and temporal scales. Only relying on scientists and concerned institutions to gather data will be the limiting factor when a lot of data is needed (Hunt et al., 2017). Utilising the help of the general public through crowdsourcing might be a solution to this problem. The help of citizen scientists has been successfully used in the past. Successful examples show how crowdsourcing data can help to better understand the subject of interest on a much wider scale than the scope and resources of a normal research project would have allowed. Examples for the successful use of crowdsourcing include *Flukebook*, a website where anyone can submit photos of whale flukes for identification. The website now has over 2 million entries which also feed into a scientific database for monitoring whales at a global scale (Levenson et al., 2015). Another example is the smartphone app *Cicada Safari* where since 2019 more than 150,000 citizen scientists have uploaded geotagged photos of cicadas, a superfamily of insects, which helped the study of re-emergence of these insects by scientists (Graber-Stiehl, 2021). The two examples highlighted above demonstrate the willingness of the general population to help gather information for analysis when the project is well conceptualised. Our method to detect spawning grounds gives the opportunity to make use of crowdsourcing to gather images of spawning grounds. The images for this study were taken using a commercially available drone at a height of 50 m during specific weather conditions to minimise reflectance that could interfere with classification. In principle an amateur drone pilot should be able to accomplish this without complicated technical guidance. The ubiquitous existence of UAVs, not least in a popular tourist destination like Iceland, provides a solid basis for crowdsourcing more data to further assess the applicability of our method and to extent the observations from our case study areas to other freshwater systems.

For further monitoring purposes, a framework building on the capabilities of Geographic Information Systems (GIS) will need to be developed to create a spatial database from the remote sensing analyses. This provides the basis for establishing change detection procedures that help to assess the spatial changes in spawning grounds on a temporal scale. Additional geographical observations from the shorelines along the lake are also derived from the remote sensing analysis as basic indicators of the different endmember classes (*e*.*g*., aquatic vegetation, vegetation, shoreline). This information provides additional useful information for a GIS database which will be valuable for geomorphological mapping of the study sites to further understand the ecosystem and see how its geography is changing beyond the nature of the nesting sites.

## Conclusions

Climate change and other anthropogenic stressors are impacting freshwater ecosystems and habitats that are within them. Salmonids are especially vulnerable during their embryonic development when highly oxygenated water and stable low temperatures are crucial. Mapping and monitoring spawning grounds can offer insights into the impacts of climate change and other stressors on the habitat of salmonid species and the freshwater ecosystem as a whole. This study shows the suitability of using low-cost UAV and pixel based classification approaches in remote sensing to detect salmonid redds in contrasting environments. The developed method has potential to be scaled to other freshwater ecosystems and salmonid species to assess the impact of anthropogenic stressors on a wider scale and to develop monitoring procedures that enable the long term observation of the changes that occur in these highly fragile environments.

