## Supplementary tables for "Monitoring fish spawning sites in freshwater ecosystems using low-cost UAV data: A case study of salmonids in lakes in Iceland"

*Table S1 Basic technical specifications of the true colour camera and battery carried by the DJI Mavic Pro drone.*

| DJI Characteristics | Specifications |
|---|---|
| Sensor | 1/2.3” (CMOS)<br>Effective pixels: 12.35 M (Total pixels: 12.71 M) |
| Sensor size | 6.3 mm x 4.7 mm |
| Lens model | FOV 78.8˚ 26 mm (35 mm format equivalent) f/2.2<br>Distortion <1.5%, focus from 0.5 m to ∞. |
| Focal length | 3.6 mm |
| Pixel pitch | 1.57 µm |
| Image size | 4000 x 3000 pixels |
| Battery capacity | 3830 mAh |
| Battery type | LiPo 3S |
| Voltage | 11.4 V |
| Overall flight time | 21 minutes |

*Table S2 Results of accuracy assessment of the two classification algorithms in Thingvallavatn. Producer’s Accuracy (PA) and User’s Accuracy (UA) by class and overall accuracy before and after post-classification improvements.*

| Class | Thingvallavatn | | | | Ellidavatn | | | |
|---|---|---|---|---|---|---|---|---|
| | Before post-classification methods | | After post-classification methods | | Before post-classification methods | | After post-classification methods | |
| | PA (%) | UA (%) | PA (%) | UA (%) | PA (%) | UA (%) | PA (%) | UA (%) |
| Spawning redds | 88.62 | 94.86 | 90.68 | 97.07 | 87.87 | 87.33 | 88.50 | 87.67 |
| Vegetation | 97.32 | 98.46 | 99.52 | 98.13 | 97.31 | 81.65 | 97.89 | 82.39 |
| Underwater rocks | 90.18 | 77.52 | 92.28 | 79.33 | 82.83 | 73.23 | 85.99 | 75.66 |
| Deep water | 98.08 | 94.50 | 99.79 | 95.57 | 94.14 | 78.95 | 96.63 | 79.43 |
| Shoreline | 84.39 | 95.23 | 85.36 | 98.13 | 42.17 | 94.05 | 41.85 | 95.29 |
| Surface rocks | 82.21 | 80.11 | 85.86 | 84.37 | 13.26 | 53.04 | 9.62 | 62.68 |
| Overall accuracy (%) | 91.27 | | 93.40 | | 78.98 | | 80.24 | |
| Kappa coefficient | 0.89 | | 0.92 | | 0.71 | | 0.72 | |
